# A chronic *Pseudomonas aeruginosa* mouse lung infection modeling the pathophysiology and inflammation of human cystic fibrosis

**DOI:** 10.1101/2024.10.07.617039

**Authors:** Mylene Vaillancourt, Diane Aguilar, Sheryl E. Fernandes, Peter A. Jorth

**Affiliations:** Department of Pathology and Laboratory Medicine, Cedars-Sinai Medical Center, Los Angeles, California, USA; Department of Biomedical Sciences, Cedars-Sinai Medical Center, Los Angeles, California, USA; Department of Medicine, Cedars-Sinai Medical Center, Los Angeles, California, USA

## Abstract

Investigation of chronic cystic fibrosis (CF) lung infections has been limited by a lack of murine models that reproduce obstructive lung pathology, chronicity of bacterial infections, and complex inflammation in human CF lung pathology. Three different approaches have been used separately to address these limitations, including using transgenic *Scnn1b-Tg* mice overexpressing a lung epithelial sodium channel to mimic the mucus-rich and hyperinflammatory CF lung environment, using synthetic CF sputum medium (SCFM) in an acute infection to induce bacterial phenotypes consistent with human CF, or using agar beads to promote chronic infections. Here, we combine these three models to establish a chronic *Pseudomonas aeruginosa* lung infection model using SCFM agar beads and *Scnn1b-*Tg mice (SCFM-Tg-mice) to recapitulate nutrients, mucus, and inflammation characteristic of the human CF lung environment. Like people with CF, SCFM-Tg-mice failed to clear bacterial infections. Lung function measurements showed that infected SCFM-Tg-mice had decreased inspiratory capacity and compliance, elevated airway resistance, and significantly reduced FVC and FEV0.1. Using spectral flow cytometry and multiplex cytokine arrays we show that, like people with CF, SCFM-Tg-mice developed inflammation characterized by eosinophil infiltration and Th2 lymphocytic cytokine responses. Chronically infected SCFM-Tg-mice developed an exacerbated mix of innate and Th1, Th2, and Th17-mediated inflammation, causing higher lung cellular damage, and elevated numbers of unusual Siglec F^+^ neutrophils. Thus, SCFM-Tg-mice represents a powerful tool to investigate bacterial pathogenesis and potential treatments for chronic CF lung infections and reveal a potential role for Siglec F^+^ neutrophils in CF inflammation.

**Importance:** Host-pathogen interaction studies of *Pseudomonas aeruginosa* cystic fibrosis (CF) lung infections have been hampered by limitations of mouse infection models. Here we combine synthetic CF sputum medium (SCFM) agar beads and *Scnn1b*-Tg transgenic mice to model the mucus obstructed airways and complex inflammatory characteristic of the human cystic fibrosis lung environment. In this model, which we name SCFM-Tg-mice, we use SCFM to cause changes in bacterial gene expression consistent with sputum collected from people with CF and the *Scnn1b-Tg* mice produce excessive airway mucus like people with CF. We show that SCMF-Tg-mice infected with *P. aeruginosa* have defects in lung function and increased inflammation that is consistent with human CF lung infections. This model can be adapted for other bacterial species and can be used to test hypotheses about bacterial pathogenesis and potential treatments in a CF human-like system.

## Introduction

*Pseudomonas aeruginosa* is a Gram-negative opportunistic bacterium responsible for persistent lung infections in people with muco-obstructive and chronic inflammation like cystic fibrosis (CF). Although primary *P. aeruginosa* infection does not seem to cause declined lung function in CF patients (*1*), adaptation and changes in *P. aeruginosa* virulence and antibiotic resistance during chronic infections are thought to be the most common cause of pulmonary exacerbations (*2*). Pulmonary exacerbations caused by bacterial infections are characterized by increased mucus production and an amplified inflammatory response leading to irreversible airway damage and a decrease in respiratory spirometry (*2, 3*). Studying bacterial interactions with the host environment has been challenging since *P. aeruginosa* modulates its gene expression in response to environmental nutrients and stress conditions, making *in vitro* characterization poorly relevant to *in vivo* infections (*4, 5*). To overcome this issue, a synthetic CF sputum-mimicking medium (SCFM2) was developed, and *P. aeruginosa* grown in SCFM2 have genetic fitness determinants and gene expression profiles that mirror bacteria grown in sputum collected from people with CF (*6–8*). However, even with SCFM2, *in vitro* studies lack crucial host factors mediating host-pathogen interactions, including the highly inflammatory and oxidative environment produced by immune cells.

Multiple *in vivo* murine models of chronic *P. aeruginosa* infections have been developed over a period of time spanning more than 3 decades (*9*). To establish chronic *P. aeruginosa* infections in mice, different strategies have been developed such as growing the strains in aggregates or adding alginate to promote biofilm-like phenotypes (*10–12*), using fibrinogen plug models (*13, 14*), or by embedding bacteria in agar beads (*15–18*). Using these strategies, most studies were successful in establishing chronic infections in mice. However, these models do not recapitulate CF infections as they utilized healthy mice lacking key pathological lung characteristics seen in human CF disease, including mucus plug and immune cell infiltration (*11, 13, 16, 18, 19*). Cystic fibrosis transmembrane conductance regulator (CFTR) mutant mice showed higher sensitivity to *P. aeruginosa* infections and developed higher inflammation compared to WT counterparts during chronic infections (*15, 17*). However, like healthy mice, they did not spontaneously develop mucus plugs and complex inflammation underlying CF disease, and thus CFTR mutant mice are not ideal models for studying *P. aeruginosa* behavior during chronic CF infections. Notably, the inflammatory response of these models was mainly neutrophilic, while human CF lung inflammation is also characterized by eosinophilia and lymphocytosis (*20–22*). In an attempt to improve the CF mouse model, CF mice with S489X *CFTR* mutations were infected with a mucoid clinical isolate of *P. aeruginosa* embedded in tryptic soy broth agarose beads. In this agar bead-CF mouse model, CF mice suffered higher mortality than normal mice, had higher inflammation, and experienced greater weight loss, but the CF mice did not have higher bacterial burdens and also lacked the mucus plugging and pulmonary disease in people with CF (*23*). *Scnn1b* transgenic (*Scnn1b*-Tg) mice overexpress the βENaC epithelium sodium channel in their lungs, causing CF-like lung pathology including mucus accumulation and neutrophil infiltration (*24, 25*). These mice were described to spontaneously develop a juvenile asthmatic inflammation that partially resolved in early adulthood (*26*). Furthermore, these mice were shown to be more sensitive to infection and develop higher inflammatory responses in early days after *P. aeruginosa* infection (*12, 13*), making them a compelling model to study chronic bacterial infections. Most recently, SCFM2 was used to pre-culture *P. aeruginosa* prior to acute lung infection of WT C57BL/6J mice and transcriptomic analyses of bacteria in infected mice revealed that this pre-culture condition promoted improved CF gene expression phenotypes in the infecting *P. aeruginosa* compared to bacteria pre-grown on Pseudomonas Isolation Agar (*27*). Yet, the authors acknowledged several limitations and suggested that the *Scnn1b*-Tg mouse model could further recapitulate CF disease physiology, which we test here. The different mouse models were only partially characterized in terms of lung mechanics and immune response during lung infection, leaving a blind spot in our knowledge of the host-pathogen interaction during chronic infections with *P. aeruginosa*.

Here, we sought to overcome limitations of previous murine *P. aeruginosa* chronic lung infection models. First, we used *Scnn1b*-Tg mice to recapitulate the underlying inflammation and obstructive lung pathology seen in human CF disease. Next, we used agar beads to promote biofilm aggregate formation and promote chronic infection. Third, we used SCFM2 agar to recapitulate the human CF nutrient environment and promote CF-like gene expression in *P. aeruginosa*. Using this multifaceted model named SCFM-Tg-mice, we showed that SCFM-Tg-mice failed to efficiently clear bacterial infection. We used highly translational lung function measurements to demonstrate the underlying obstructive disease in these mice, and the efficiency of these parameters to track lung decline during chronic infections. We also deciphered the complex immune and inflammatory responses of this model compared to their WT littermates using spectral flow cytometry and a multiplex cytokine array. Like people with CF, chronically infected *Scnn1b*-Tg mice developed an exacerbated and complex mix of innate and Th1, Th2, and Th17-mediated inflammation, causing higher lung cellular damage. Finally, we unveiled new potential players in this complex inflammatory response.

## Results

### Bacterial clearing is impaired in SCFM-Tg-mouse model

To establish a chronic infection in mice, we embedded *P. aeruginosa* PAO1 in synthetic CF sputum medium (SCFM2)(*7*) agar beads (Fig. 1, A and B). We intratracheally inoculated 1×10^6^ colony-forming units (CFUs) or sterile SCFM2 agar beads into *Scnn1b*-Tg mice or their wild-type (WT) littermates. After 7 days of infection, the bacterial load was more than 5-fold higher in *Scnn1b*-Tg compared to their WT littermates (Fig. 1C). These results confirm that *Scnn1b*-Tg mice have impaired bacterial clearing during chronic infection.

**Fig. 1.**
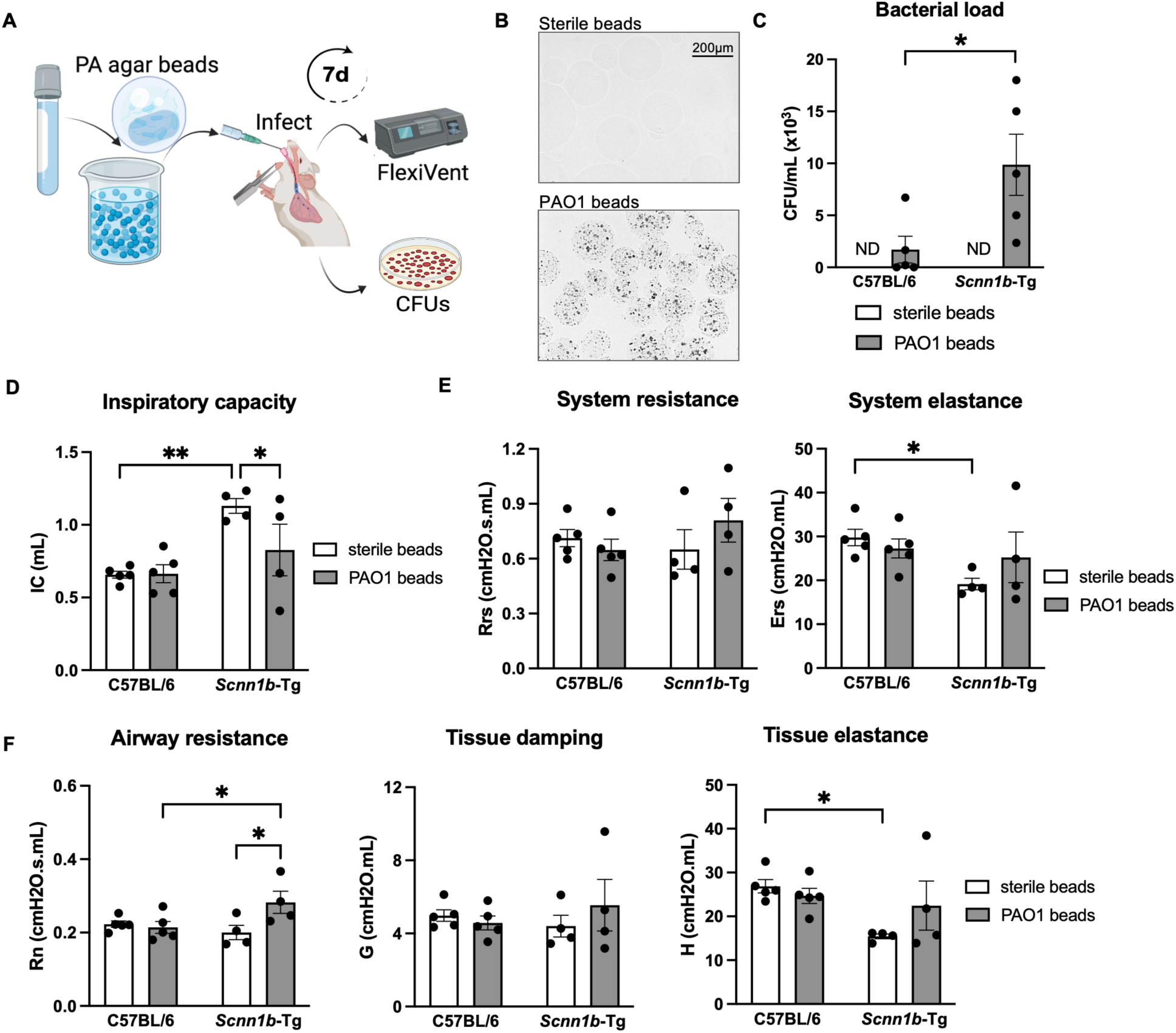
Bacterial clearance is impaired in *Scnn1b*-Tg mice and increases airway resistance during chronic infection. (**A**) WT C57BL/6 or *Scnn1b*-Tg mice were intratracheally inoculated with sterile or 1×10^6^ CFU PAO1-laden SCFM2-agar beads for 7 days. (**B**) Representative microscopic images of sterile and PAO1-laden SCFM2-agar beads. (**C**) Bacterial load 7 days post-infection. Bacterial load was determined by CFU/mL. (**D-F**) Lung function measurements obtained using the flexiVent (SCIREQ). (**D**) Inspiratory capacity using a deep inflation technique. (**E**) System resistance and elastance parameters acquired by the single frequency forced oscillation maneuver. (**F**) Airway resistance, tissue resistance (damping) and elastance obtained from the low frequency forced oscillation technique. n=4-5 mice/group. \**p*<0.05, ** *p*<0.01. See Table S1 for statistical tests used and exact *p*-values.

### SCFM-Tg-mice develop a mixed obstructive and restrictive lung disease

To determine the lung function after 7 days of infection, mice underwent measurement of respiratory mechanics with the flexiVent (SCIREQ). At baseline, *Scnn1b*-Tg mice had significantly higher inspiratory capacities compared to their WT littermates (Fig. 1D and fig. S1A) accompanied by a lower system elastance (Fig. 1E and fig. S1B). This decreased elastance was caused by a loss in lung tissue resistance and elasticity without any difference in the central airway resistance (Newtonian resistance) (Fig. 1F and fig. S1C). Chronic infection significantly decreased the inspiratory capacity of *Scnn1b*-Tg mice (Fig. 1D) and increased airway resistance (Fig. 1F). The pressure-volume (PV) curves highlighted higher static compliance and an increased area of hysteresis (Fig. 2, A and B, and fig. S1D) in *Scnn1b*-Tg mice at baseline, confirming the presence of emphysema. Both compliance and hysteresis were decreased during chronic infection, indicating restriction during inhalation (Fig. 2, A and B). The negative pressure-driven forced expiration (NPFE) maneuver (Fig. 2, C and D, and fig. S1F) showed an increase in the forced vital capacity (FVC), forced expiratory volume within 0.1 s (FEV0.1), and forced expiratory flow at 0.1 s (FEF0.1) in *Scnn1b*-Tg mice compared to WT mice at baseline, consistent with an obstructive pathology. All three parameters were significantly decreased in the SCMF-Tg-mice chronic infection. The decreased FEV0.1/FVC ratio seen in SCFM-Tg-mice (68±3 vs 87±3 in WT mice) supports reduced lung capacity in this model. Finally, a significant interaction was detected between the infection with *P. aeruginosa* and the *Scnn1b*-Tg genotype on the airway resistance, the FEV0.1, and the peak expiratory flow (PEF) (Table S1). This suggests a higher sensitivity to lung function decline in *Scnn1b*-Tg mice during chronic infection with *P. aeruginosa*. Taken together, the lung mechanics confirmed an initial obstructive lung disease in *Scnn1b*-Tg mice and demonstrated the establishment of a mixed obstructive/restrictive pathology during chronic infection in the SCFM-Tg-mice.

**Fig. 2.**
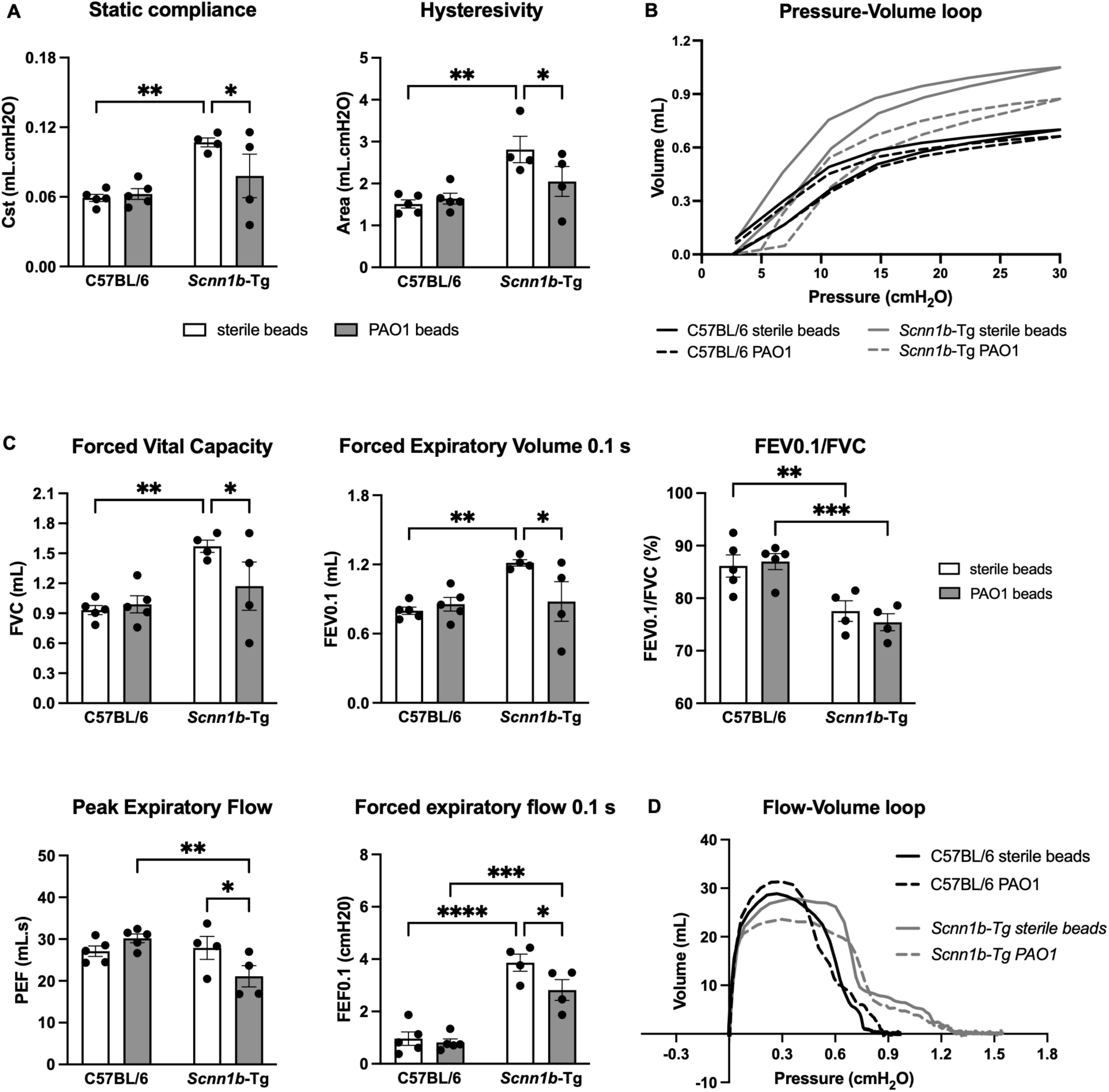
*Scnn1b*-Tg mice develop mixed obstructive and restrictive lung disease during chronic infection. WT C57BL/6 or *Scnn1b*-Tg mice were intratracheally inoculated with sterile or 1×10^6^ CFU PAO1-laden SCFM2-agar beads for 7 days. (**A-D**) Lung function measurements obtained using the flexiVent (SCIREQ). (**A**) Static compliance and hysteresivity obtained by a pressure-volume (PV) loop. (**B**) Representative image of PV-loop. (**C**) FVC, FEV0.1, FEV0.1/FVC, PEF and FEF0.1 obtained from the forced expiratory volume perturbation. (**D**) Representative image of the forced expiratory volume perturbation. n=4-5 mice/group. \**p*<0.05, \*\**p*<0.01, \*\*\**p*<0.001, \*\*\*\**p*<0.0001. See Table S1 for statistical tests used and exact *p*-values.

### Atypical neutrophils are increased in the SCFM-Tg model

*Scnn1b*-Tg mice were previously described to develop chronic airway inflammation characterized by increased macrophages, neutrophils, eosinophils, and lymphocytes (*24, 25*). We confirmed the presence of lung inflammation in the bronchoalveolar lavage (BAL) of uninfected *Scnn1b*-Tg mice (figs. S2 and S3). To determine whether this underlying inflammation could modify the inflammatory response to bacterial infection, we infected *Scnn1b*-Tg mice or their WT littermates with PAO1-laden or sterile SCFM2 agar beads for 7 days (Fig. 3A). We then performed an inflammatory flow cytometry panel on the total lung and analyzed the cells by spectral flow cytometry (Fig. 3B). As expected, total inflammatory cells were increased in infected mice of both genotypes (Fig. 4A). Alveolar and monocyte-derived macrophages were also increased in infected mice (Fig. 4, B and C). Infected *Scnn1b*-Tg mice showed a mild increase in classical monocytes (Fig. 4D), while the inflammation in WT mice was characterized by other CD11b+ myeloid cells (Fig. 4E). Although eosinophils were higher in BAL of uninfected *Scnn1b*-Tg compared to WT mice (Fig. S2), they were not further increased during chronic infection (Fig. 4F). A surprising finding in infected *Scnn1b*-Tg mice was the upregulation of an atypical Siglec-F^+^ neutrophil subset (Fig. 4, H and I), despite no difference in total neutrophils between the two mouse genotypes.

**Fig. 3.**
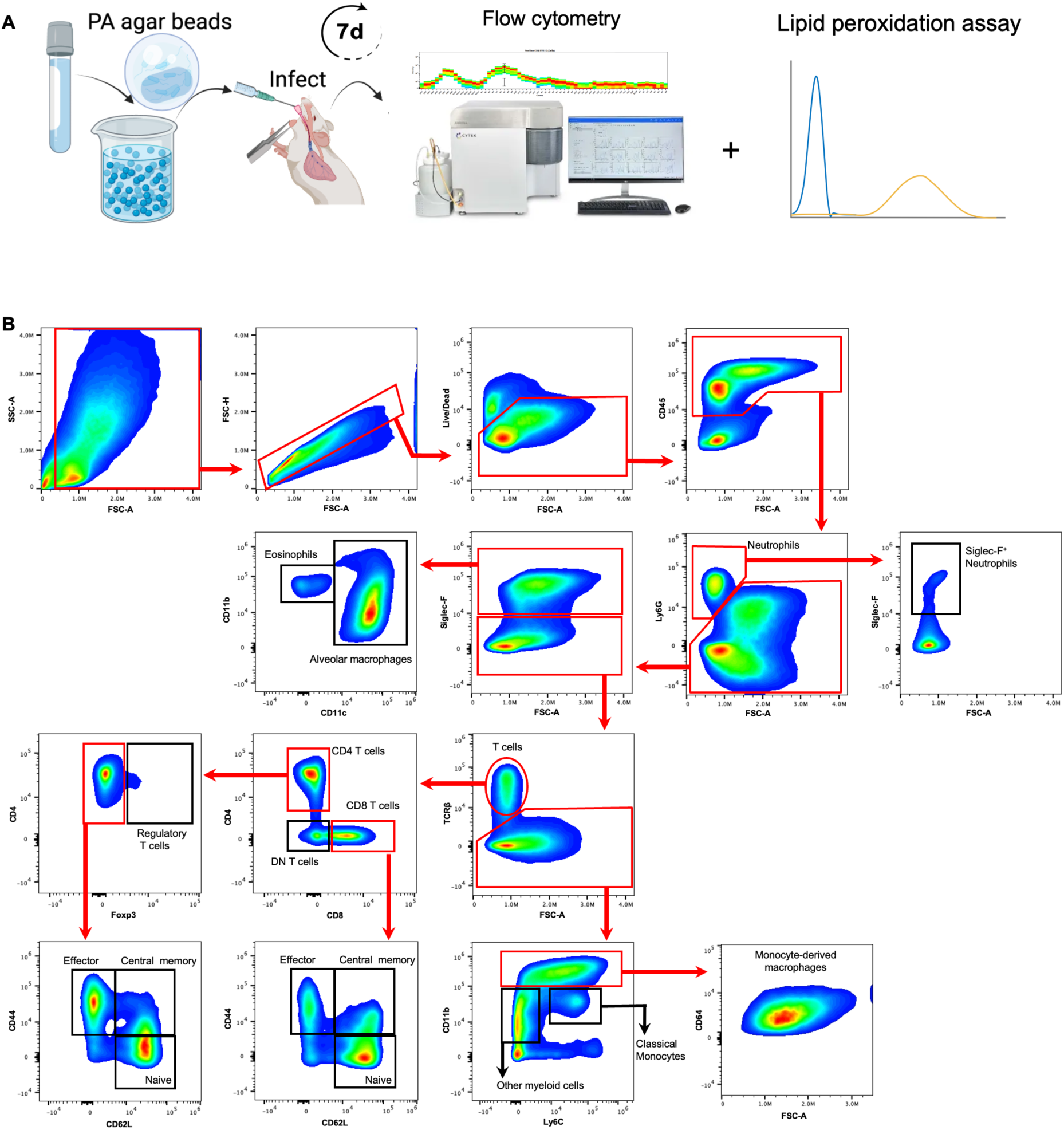
Chronic infection in WT C57BL/6 or *Scnn1b*-Tg mice for immune response by flow cytometry. (**A**)WT C57BL/6 or *Scnn1b*-Tg mice were intratracheally inoculated with sterile or 1×10^6^ CFU PAO1-laden SCFM2-agar beads for 7 days. (**B**) Gating strategy used to identify immune cell response during chronic infection. Cells were isolated from enzymatically digested mouse lungs, and, after the exclusion of doublets and debris, live and immune cells were identified by LIVE/DEAD staining and CD45 staining. Neutrophils (Ly6G^+^) were isolated and gated for Siglec F marker. Then, Ly6G^-^ and Siglec F^+^ cells were selected to differentiate alveolar macrophages (Siglec-F^+^, CD11c^+^) and eosinophils (Siglec-F^+^, CD11b^+^, CD11c^-^). T cells (TCRβ^+^) were then separated from the rest of Sigle F^-^ cells. CD4^+^ and CD8^+^ were separated from the double-negative (DN) subset. CD4^+^ and Foxp3^+^ cells were isolated, while Foxp3^-^ cells were separated by the CD44 and CD62L markers to identify naïve CD4^+^ T cells (CD44^-^, CD62L^+^), effector CD4^+^ T cells (CD44^+^, CD62L^-^), and central memory CD4^+^ T cells (CD44^+^, CD62L^+^). CD8^+^ T cells were also separated with the same markers CD44+ and CD62L. Finally, TCR^-^ cells were further separated using Ly6C and CD11b markers to identify monocyte-derived macrophages (CD11b^High^, Ly6C^+/-^, CD64^+^, FSC-A^high^), classical monocytes (CD11b^+^, Ly6C^+^), and other myeloid-derived cells (CD11b^+^, Ly6C^-^).

**Fig. 4.**
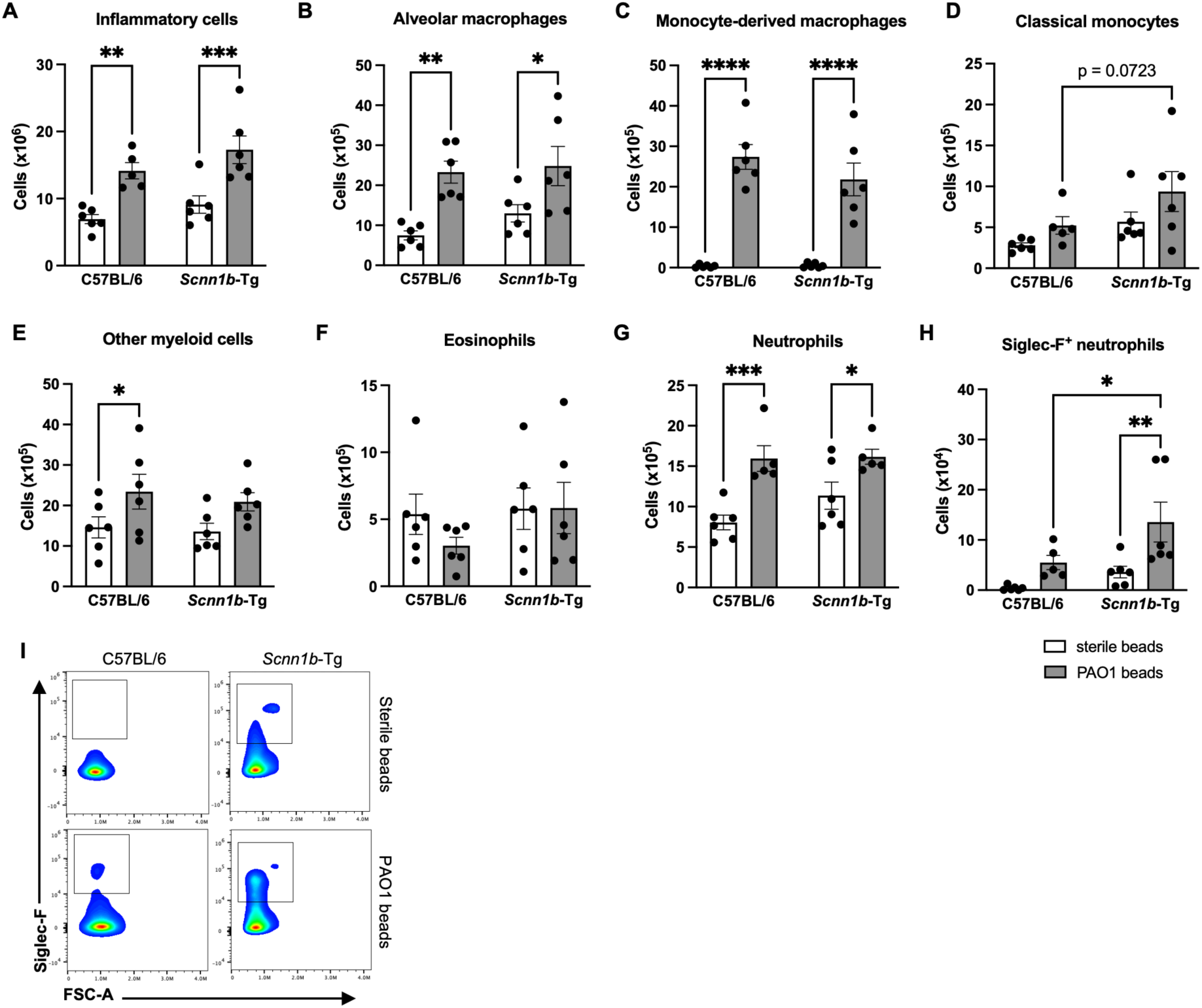
*Scnn1b*-Tg mice lung inflammation is characterized by an increase in atypical neutrophils. (**A**) Inflammatory cells were increased in both *Scnn1b*-Tg mice and their WT littermates. (**B-E**) Different innate cells were upregulated in both genotype during chronic infection. (**B**) Alveolar macrophages. (**C**) Monocyte-derived macrophages. (**D**) Classical monocytes. **E.** Other myeloid cells. (**F**) Eosinophils were not upregulated during chronic infection with *P. aeruginosa*. (**G**) Neutrophils were upregulated during chronic infection but not modulated by the genotype. (**H-I**) An atypical Siglec F^+^ neutrophil subset was upregulated in *Scnn1b*-Tg mice during chronic infection. n=6 mice/group \**p*<0.05, \*\**p*<0.01, \*\*\**p*<0.001, \*\*\*\**p*<0.0001. See Table S1 for statistical tests used and exact *p*-values.

### Effector lymphocytes are a characteristic of *Scnn1b*-Tg immune response

To further characterize the inflammatory response of *Scnn1b*-Tg mice during chronic infection, we looked at different lymphocyte subtypes and their activation state. As expected during chronic infection, infiltrating T cells were present in the lung tissues of all infected mice (Fig. 5A). Both CD4^+^ and CD8^+^ T cells were significantly increased in *Scnn1b*-Tg compared to WT mice (Fig. 5, B and C). We then separated the cells on whether they were activated (CD44^+^, CD62L^-^) or naïve (CD44^-^, CD62L^+^). In each subtype, a small proportion of cells were positive for both markers (CD44^+^, CD62L^+^) and were qualified as central memory T cell (*28*). Infected mice of both genotypes showed increased effector CD4^+^ T cells during chronic infection (Fig. 5D), but effector CD4^+^ T cells were also significantly higher in infected *Scnn1b*-Tg mice compared to WT mice. No change in naïve CD4^+^ T cells was observed during chronic infection (Fig. 5D). We also observed a mild but non-significant increase in central memory T cells for both WT and *Scnn1b*-Tg mice (Fig. 5D). Effector and central memory CD8^+^ T cells were both increased during infection, but not modulated by the mouse genotypes (Fig. 5E). Naïve CD8^+^ T cells increased in *Scnn1b*-Tg mice during infection and were significantly higher than in WT mice (Fig. 5E). Because regulatory T cells can modulate the immune response (*29*), we looked at whether they were differentially present in the infected lungs of WT and *Scnn1b*-Tg mice. Although regulatory T cells were more abundant in infected mice, there was no difference between the two genotypes (Fig. 5F). Finally, we observed a significant increase in CD4^-^ CD8^-^ double negative (DN) T cells in the infected lungs of *Scnn1b*-Tg mice (Fig. 5G).

**Fig. 5.**
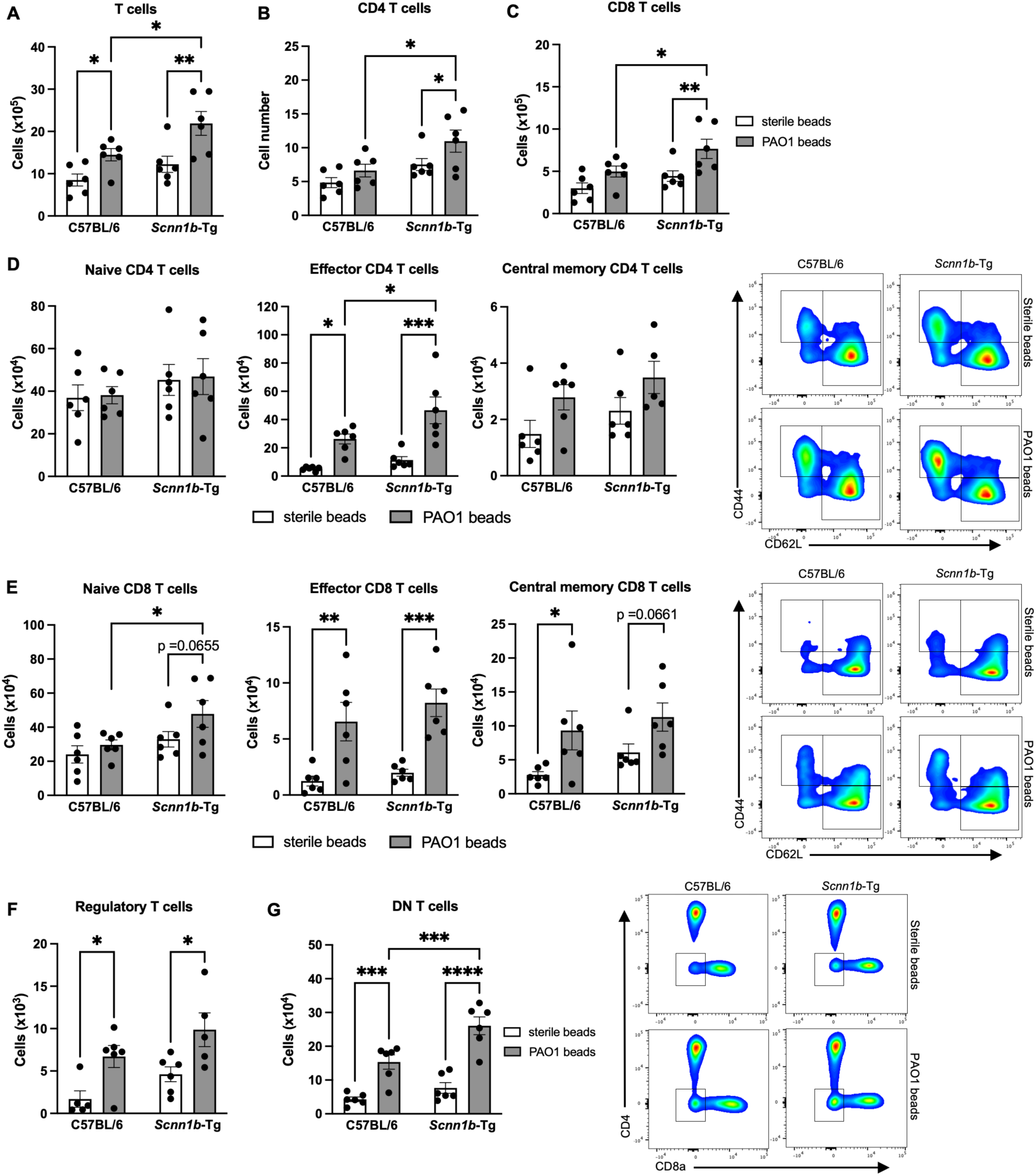
*Scnn1b*-Tg immune response is characterized by effector T cells. (**A**) Total T cells were significantly increased during chronic infection and even more in the *Scnn1b*-Tg mice. (**B-C**) This increase in T cells was explained by higher numbers of CD4^+^ (**B**) and CD8^+^ (**C**) T cells. (**D**) Activation state of CD4^+^ T cells. No difference was seen in naïve CD4^+^ T cells. During chronic infection, a significant upregulation of effector T cell was observed in both genotypes and this increase was greater in *Scnn1b*-Tg mice compared to their WT littermates. A modest but non-significant increase was detected for central memory T cells in infected mice. (**E**) Activation state of CD8^+^ T cells. During chronic infection, a significant increase in naïve CD8^+^ T cell was observed in *Scnn1b*-Tg mice. Effector T cells were also increased in both genotypes. A modest increase of central memory CD8 was detected for both genotypes. (**F**) Regulatory T cells were also increased in all infected mice but not modulated by the genotype. **G.** Double-negative (DN) cells were significantly increased in all infected mice and were significantly higher in *Scnn1b*-Tg mice compared to their WT littermates. n=6 mice/group \**p*<0.05, \*\**p*<0.01, \*\*\**p*<0.001, \*\*\*\**p*<0.0001. See Table S1 for statistical tests used and exact *p*-values.

### *Scnn1b*-Tg mice develop exacerbated innate inflammation during chronic infection

Next, we performed a 29-plex cytokine array on the whole mouse lungs to quantify inflammatory signaling. As expected, pro-inflammatory cytokines like IL-6, IL-1β, and TNFα were upregulated during infection (Fig. 6, A and B). The monocyte/macrophage chemokines MIP-1α and CXCL10, as well as the neutrophil chemokines KC/GRO and MIP-2 were also upregulated and significantly higher in *Scnn1b*-Tg compared to WT mice (Fig. 6, A, C, and D). Furthermore, there was a significant interaction between the infection with *P. aeruginosa* and the *Scnn1b*-Tg genotype on KC/GRO levels (Table S1), demonstrating a synergistic effect of these parameters on the neutrophil-attractant chemokine. The increased inflammatory cytokines and chemokines in *Scnn1b*-Tg mice are surprising since the counts for most of the cell types of the innate response were not different between infected *Scnn1b*-Tg and WT mice (Fig. 4). This could mean that although the cell numbers are similar, innate cells in *Scnn1b*-Tg mice are hyperactivated during infection. However, whether hyperactivation is due to differences in bacterial loads, or in the immune cells is yet to be determined. We also looked at how T cell cytokines were modulated during chronic infection (Fig. 6, A and E). MIP-3α, a chemokine expressed by activated macrophages and a strong chemoattractant for lymphocytes (*30*), was increased in infected mice and significantly higher in *Scnn1b*-Tg mice. IL-15, a promotor of CD8^+^ T cell proliferation (*31*), was only increased in C57BL/6 mice. IL-16, a major CD4^+^ T cell activator (*32*), was highly expressed and significantly increased in infected *Scnn1b*-Tg mice. The increase of MIP-3α and IL-16 may explain the high numbers of effector T cells seen in *Scnn1b*-Tg mice (Fig. 5).

**Fig. 6.**
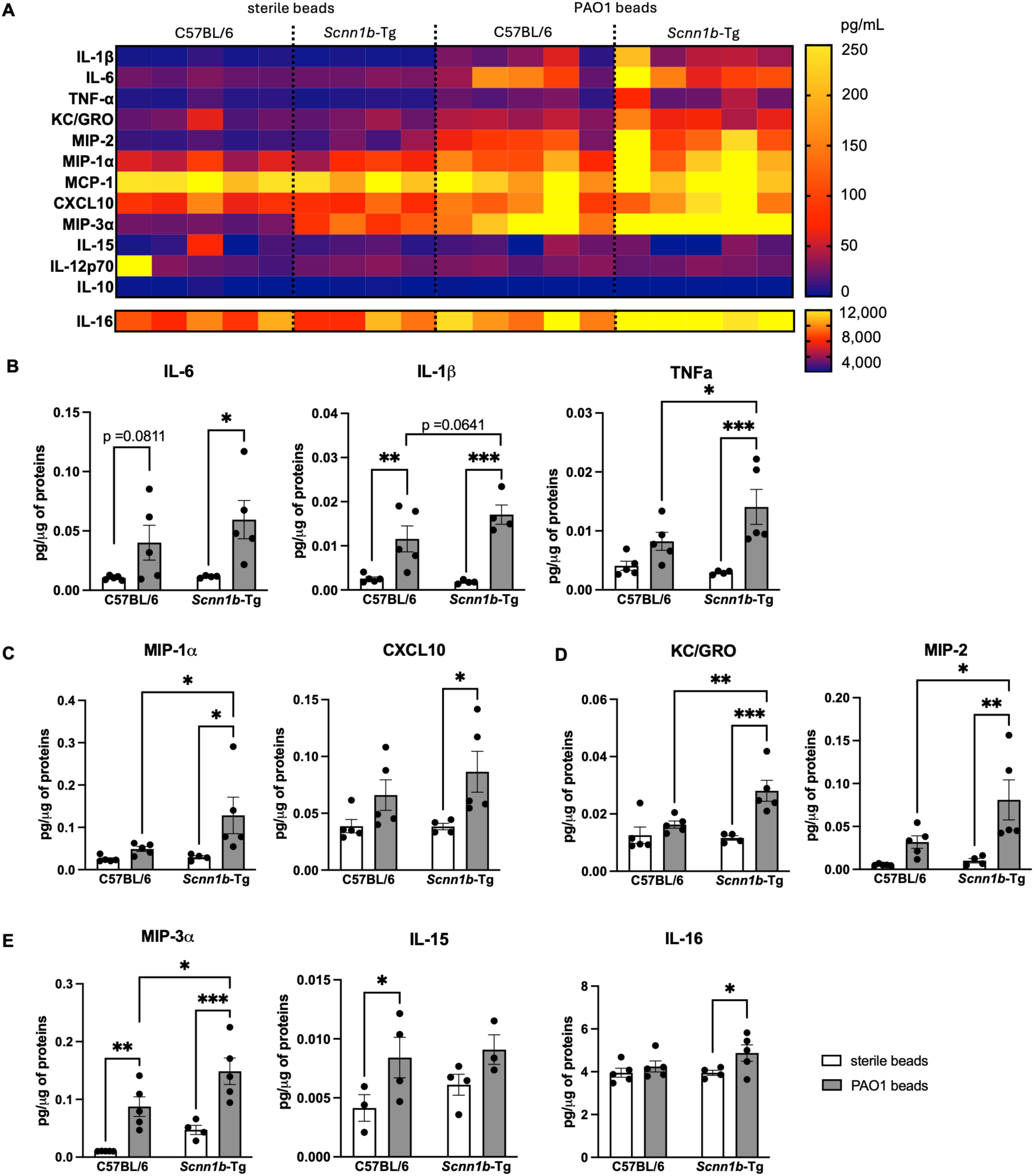
*Scnn1b*-Tg mice develop exacerbated innate inflammation during chronic infection. (**A**) Quantification (pg/μg of proteins) of pro- and anti-inflammatory cytokines and chemokines in whole lung lysates of chronically infected mice. (**B**) Inflammatory cytokines IL-6, IL-1β, and TNF-α were upregulated in all infected mice. IL-1β and TNF-α levels were higher in *Scnn1b*-Tg mice compared to their WT littermates. (**C**) Monocytes/macrophages chemoattractant MIP-1α and CXCL10 were significantly upregulated in infected *Scnn1b*-Tg mice. (**D**) Neutrophil chemoattractants were significantly upregulated in infected *Scnn1b*-Tg mice compared to their WT littermates. (**E**) Lymphocyte chemoattractant MIP-3α was upregulated in all infected mice but was higher in *Scnn1b*-Tg mice. IL-15 was increased in infected WT C57BL/6 mice only, while and IL-16 was only upregulated in *Scnn1b*-Tg mice. n=4-5 mice/group \**p*<0.05, \*\**p*<0.01, \*\*\**p*<0.001. See Table S1 for statistical tests used and exact *p*-values.

### Chronic infection leads to dysfunctional lymphoid-mediated inflammation in *Scnn1b*-Tg mice

To further understand the type of inflammation present during *P. aeruginosa* chronic infection in our model, we looked at levels of typical cytokines present during different types of inflammation. Type 1 inflammation is driven by Th1 lymphocytes and triggered in response to harmful pathogens or injury (*33*). IL-2, which promotes the survival and differentiation of naïve T cells in Th1 and Th2 (*34, 35*), was upregulated during infection and increased in *Scnn1b*-Tg mice (Fig. 7, A and B). IFNγ and IL-27 are secreted during this type 1 response and were significantly upregulated in infected *Scnn1b*-Tg mice, while being moderately increased in WT mice (Fig. 7, A and B). A type 1 inflammation response to bacterial infection is expected, but these results support an exacerbated response to infection in *Scnn1b*-Tg mice. We further looked at cytokines involved in the types 2 and 3 inflammation. Type 2 inflammation is an overactive immune response and is mainly seen in asthmatic and allergic diseases (*36*), is correlated with declined lung function, and is common during *P. aeruginosa* infections in CF (*37*). Type 2 inflammation is characterized by eosinophilia and high levels of IL-4, IL-5, and IL-33 (*33, 38*). Our cytokine array revealed that IL- 4 and IL-5 levels were significantly higher in uninfected *Scnn1b*-Tg mice compared to WT mice (Fig. 7, A and C). This was consistent with the presence of eosinophils in the BAL of uninfected *Scnn1b*-Tg mice (fig. S2). IL-4 is secreted by Th2 lymphocytes, eosinophils, basophils, and mast cells, and induces differentiation of naïve helper cells into Th2 lymphocytes (*39*). On the other side, IL-5 is produced by Th2 cells and is a key mediator of eosinophil activation (*40*). These results support the presence of chronic type 2 inflammation in *Scnn1b*-Tg mice even before infection. During chronic infection, it was surprising to see that while IL-4 was downregulated, IL-5 levels did not change (Fig. 7, A and C). IL-33, another cytokine involved in the maturation of Th2 cells and activation of eosinophils (*41*), was further increased during chronic infection (Fig. 7, A and C). Although type 2 inflammation is not typically triggered during bacterial infections, these results show that it is still present and may play a role in the dysfunctional inflammatory response in *Scnn1b*-Tg mice. IL-17A is a cytokine produced by Th17 cells and plays a key role in T cell-mediated neutrophil mobilization and activation (*33, 42, 43*). IL-17-mediated inflammation was also described in CF patients and was correlated with pulmonary exacerbations and infection with *P. aeruginosa* (*44, 45*). IL-17A was upregulated in all infected mice but was significantly higher in the lungs of *Scnn1b*-Tg mice (Figure 7, A and D). Furthermore, as for KC/GRO, there was a significant interaction between the infection with *P. aeruginosa* and the *Scnn1b*-Tg genotype on IL-17A levels, demonstrating a synergistic effect of these parameters on the neutrophil-attractant chemokine (Table S1). Finally, we used z-scores to compare *Scnn1b*-Tg and WT C57BL/6 inflammation in uninfected (sterile beads) and infected mice (Fig. 7E). The high z-scores of IL-2, MIP-3α, IL-4, and IL-5 further highlighted a T cell-mediated type 2 inflammation in uninfected *Scnn1b*-Tg mice, while the type 3 inflammation (IL-17A) was the most differentially upregulated during chronic infection (Fig. 7E). These results underscore key interplays between activated T cells, eosinophils, and neutrophils during chronic *P. aeruginosa* infection and demonstrate a complex inflammatory response in our animal model similar to CF disease.

**Fig. 7.**
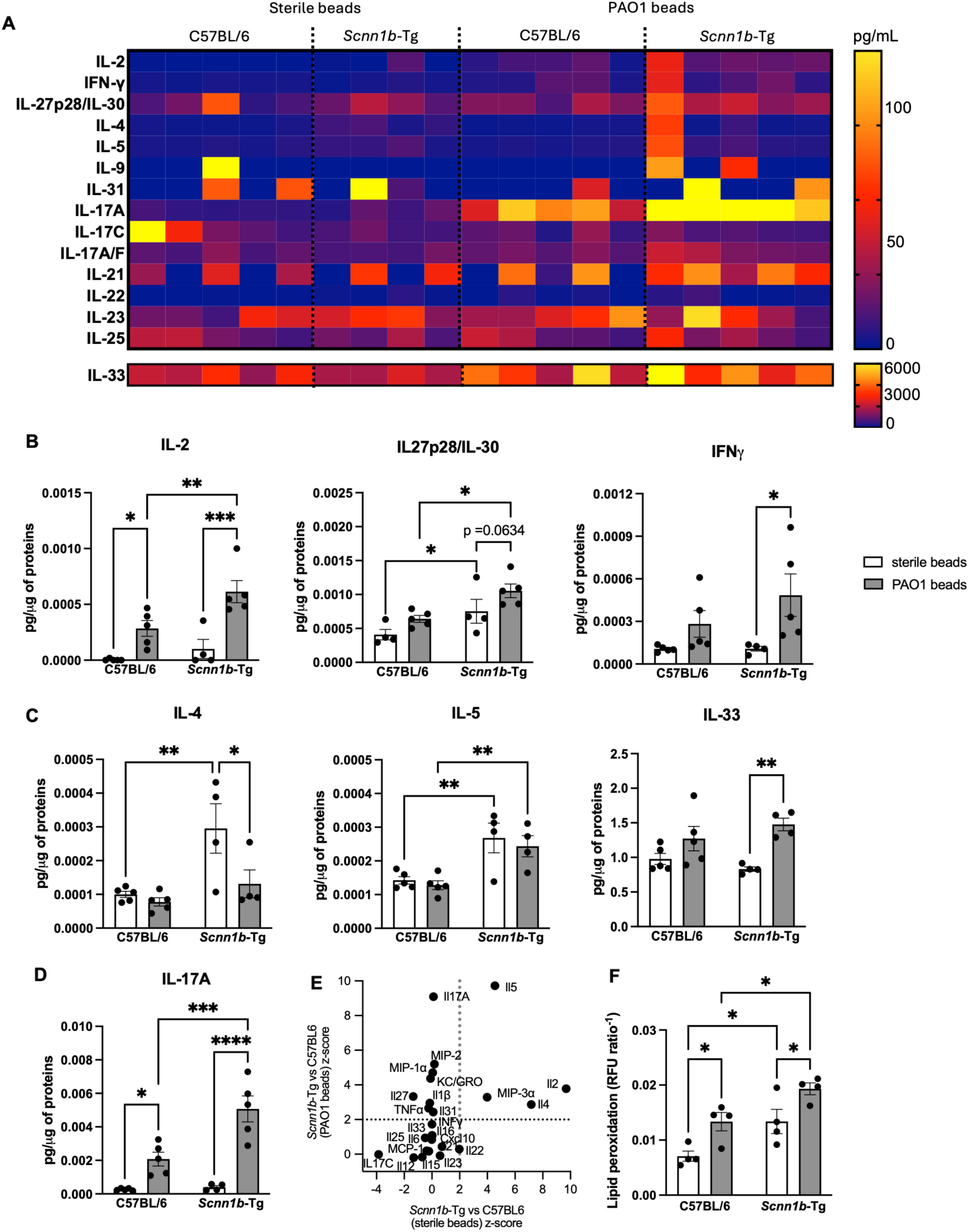
Chronic infection leads to dysfunctional lymphoid-mediated inflammation in *Scnn1b*-Tg mice. (**A**) Quantification (pg/μg of proteins) of type 1, 2 and 3 inflammation cytokines and chemokines in whole lung lysates of chronically infected mice. (**B**) Type 1 inflammation lymphokines IL-2, IL-27p/28/IL-30, and IFN-γ were significantly upregulated in infected *Scnn1b*-Tg mice. (**C**) Type 2 inflammation lymphokines IL-4 and IL-5 were upregulated in uninfected *Scnn1b*-Tg mice. Although IL-4 was downregulated during infection, IL-5 levels were maintained. IL-33 was upregulated in *Scnn1b*-Tg mice during chronic infection. (**D**) The type 3 inflammation cytokine IL-17 was upregulated in all mice and further increased in *Scnn1b*-Tg mice. (**E**) Z-scores highlight a type 2 and lymphoid inflammation in uninfected *Scnn1b*-Tg mice compared to their WT littermates. IL-17 is the most differentially upregulated cytokine in these mice during infection. (**F**) *Scnn1b*-Tg have higher lung tissue damage at baseline. Chronic infection caused increased lipid peroxidation in all infected mice but was greater in *Scnn1b*-Tg mice. n=4-5 mice/group \**p*<0.05, \*\**p*<0.01, \*\*\**p*<0.001, \*\*\*\**p*<0.0001. See Table S1 for statistical tests used and exact *p*-values.

### Chronic inflammation is associated with increased lung damage in *Scnn1b*-Tg mice

Exacerbated chronic inflammation is known to induce high oxidative stress resulting in tissue damage in lung diseases (*46*). Lipid peroxidation is a measure of cellular damage and is increased in the lungs of CF patients (*47–49*). To verify whether the high inflammatory environment induced lung tissue damage in *Scnn1b*-Tg mice, we measured lipid peroxidation on the whole lung tissue. Consistent with the cytokine assay, we found increased lipid peroxidation in uninfected lungs of *Scnn1b*-Tg mice (Fig. 7F). Chronic infection increased lipid peroxidation in both genotypes but was further increased in *Scnn1b*-Tg compared to WT mice (Fig. 7F). These results confirmed increased tissue damage at baseline and during chronic infection in our mouse model similar to what is found in CF disease.

## Discussion

Establishing a murine model of *P. aeruginosa* chronic infection that mimics the complex CF lung environment has been challenging investigators for decades (*9*). Researchers have used different engineered murine models, but most of them failed to develop spontaneous chronic infections or were limited in recapitulating key aspects of CF lung pathology (*9*). In this study, we used the *Scnn1b*-Tg mouse (*24, 25*), a model already described to develop mucus plugs, inflammation, and obstructive disease, to mimic the CF lung environment. We also used sputum-mimicking media to embed *P. aeruginosa* to further simulate the nutritional and biofilm-promoting environment found in CF airways (*7*). We assessed pulmonary function using SCIREQ flexiVent and used highly translational parameters to characterize our model (Figs. 1 and 2, and fig. S1). We showed that chronic *P. aeruginosa* infection decreased inspiratory capacity and compliance, elevated airway resistance, and significantly reduced FVC and FEV0.1, an equivalent of the gold standard FEV1 measure in clinical spirometry (*50*). We also demonstrated a greater susceptibility to lung function decline for *Scnn1b*-Tg mice compared to WT C57BL/6 littermates (Table S1). Similar to human CF disease, our model mainly develops obstructive lung disease that is mixed with restrictive disorder during chronic infection with *P. aeruginosa*.

CF inflammation is a complex mix of innate and lymphoid inflammation(*22*). It is characterized by high secretion of pro-inflammatory cytokines like IL-6, IL-1β, and TNFα, but also type 2 (IL-4, IL-5, IL-33) and type 3 (IL-17) inflammatory cytokines (*22, 44, 51*). Our characterization of the lung immune response also showed a complex inflammatory environment resembling the one described in people with CF and is summarized in Fig. 8. We first confirmed the presence of chronic inflammation at baseline *Scnn1b*-Tg mice characterized by higher counts of myeloid- and lymphoid-derived cell counts in the alveolar space (figs. S2 and S3). During chronic infection, although the innate cell counts were similar between *Scnn1b*-Tg mice and their WT littermates (Fig. 3), we showed an exacerbated inflammation by the highly secreted pro-inflammatory cytokines IL-6, IL-1β, and TNFα, and chemokines MIP-1α, CXCL10, MIP-2, and KC/GRO (Fig. 5). This suggests that monocytes, macrophages, and neutrophils may be hyperactivated in the *Scnn1b*-Tg lungs. We also describe for the first time the presence of an atypical neutrophil subset positive for the surface lectin Siglec F (Fig. 4, H and I, and fig. S2). Little is known about these Siglec F^+^ neutrophils but they were recently described as long life-span and high ROS activity neutrophils, and were shown to be deleterious in tissue fibrosis and tumor tolerance (*52–55*). In a mouse nasal mucosae infection model, high IL-17 secreting Siglec F^+^ neutrophils were associated with a better clearance *Bordetella pertussis* (*56*). In our study, it is not clear whether the high recruitment of Siglec F^+^ neutrophils is deleterious or a response to the infection to tentatively clear *P. aeruginosa.* Shin et al. also described the induction of these unique neutrophils in air pollutant-induced lung damage (*57*). Siglec F^+^ neutrophils were associated with exacerbated asthma and triggered emphysema by producing high levels of cysteinyl leukotrienes and neutrophil extracellular traps. *In vivo*, Siglec F^+^ neutrophils enhanced IL-5 and IL-13 production by Th2 cells and IL-17 secretion by CD4^+^ T cells (*57*). Here, since Siglec F^+^ neutrophils were already present in the BAL of uninfected *Scnn1b*-Tg mice and recruited during chronic infection, we hypothesize that this unique population could be involved in higher type 2 and 3 inflammatory cytokines seen in the *Scnn1b*-Tg mice during chronic infections. Presently, Siglec F^+^ neutrophils have not been described in people with CF. However, a unique neutrophil subset called low-density neutrophils was also found in the blood of CF patients and other inflammatory diseases (*58, 59*). Like Siglec F^+^ neutrophils, low-density neutrophils showed increased IL-17 production, enhanced degranulation, and decreased phagocytosis (*58, 60, 61*). Furthermore, this particular neutrophil subset was associated with pulmonary exacerbations, declined lung function, and disease progression in CF patients (*59, 62*). Although we cannot directly compare low-density neutrophils with our Siglec F^+^ neutrophils, we speculate that these two neutrophil subsets may have similar effects on chronic inflammation. Since Siglec F^+^ neutrophils, but not total neutrophil counts, were increased in *Scnn1b*-Tg mice and by the infection (Fig. 4 and fig. S2), we speculate that this specific neutrophil subset may modulate inflammation during chronic infection with *P. aeruginosa*. To our knowledge, this is the first time these Siglec F^+^ neutrophils have been observed in a CF-like model.

**Fig. 8.**
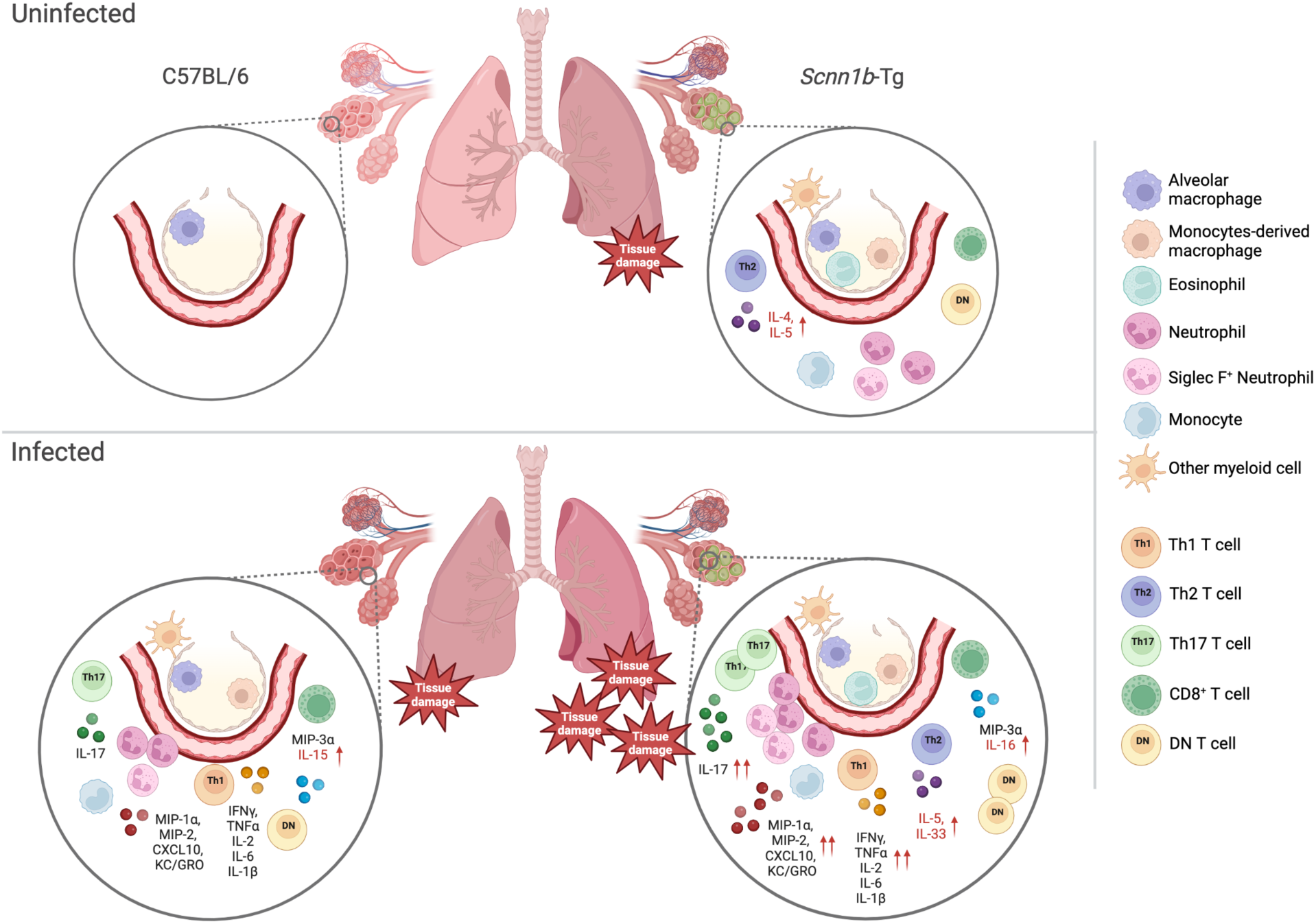
Summary of lung inflammation in C57/BL/6 and *Scnn1b*-Tg mice during chronic infection with *P. aeruginosa*. Healthy lungs from C57BL/6 mice are characterized by alveolar macrophages. Uninfected *Scnn1b*-Tg mice show underlying inflammation characterized by the presence of alveolar and monocyte-derived macrophages, monocytes, other myeloid cells, and effector T cells. Conventional and Siglec F^+^ neutrophils are also present in the BAL of uninfected *Scnn1b*-Tg mice. A type 2 inflammation, demonstrated by the presence of eosinophils and IL-4 and IL-5, is present at baseline in the *Scnn1b*-Tg lung environment. During chronic infection, both C57BL/6 and *Scnn1b-*Tg immune responses are characterized by infiltration of innate and T cells and high levels of type 1 and 3 inflammation cytokines and chemokines (IL-1β, IL-2, IL-6, IL-17, TNFα, IFNγ, MIP-1α, MIP-2, MIP-3α and KC/GRO). In addition to being exacerbated, *Scnn1b*-Tg inflammation is characterized by a sustained type 2 inflammation, a marked IL-17/neutrophil interplay and the recruitment of unconventional Siglec F^+^ neutrophils. The higher inflammation is associated with higher lung tissue damage in *Scnn1b*-Tg mice. Cytokines in red font are cytokines expressed in the specific genotypes. Red arrows indicate where the cytokine production is increased relative to the other genotype.

Another feature of our model is the evident infiltration of effector CD4^+^ T cells at baseline (Fig. 5 and fig. S3) and their proliferation during chronic infection in the *Scnn1b*-Tg mice (Fig. 5). Lymphocytosis was also described in CF patients (*22, 63*) and a skewed response toward Th2 and Th17 inflammation in these patients was associated with *P. aeruginosa* infection, declined lung function, and higher mortality (*37, 44*). Mueller *et al*. demonstrated how this skewed T cell response could result from the CFTR deficiency in lymphocytes (*64*). Interestingly, *Scnn1b*-Tg mice, which do not lack the CFTR channel, also show Th2 and Th17 responses during chronic infection (Figure 7). *P. aeruginosa* toxins can also increase type 2 inflammation resulting in higher eosinophil infiltration, cytokine (IL-4, IL-13) and IgE secretion, and mucus production (*65*). It is unclear whether *P. aeruginosa* had a role in the maintenance of the Th2 response in our *Scnn1b*-Tg mice since no Th2 response was induced in WT mice (Fig. 7). This difference could mean that either the *P. aeruginosa* secretome is modulated by the CF-like environment, or that other independent factors are responsible for the Th2 maintenance in the *Scnn1b*-Tg mice. Fritzsching *et al.* described spontaneous type 2 inflammation in juvenile *Scnn1b*-Tg mice caused by impaired mucous clearance (*26*). These mice developed an exacerbated eosinophilic response to allergen challenge. In our study, our *Scnn1b*-Tg mice also had higher baseline levels of IL-4 and IL-5 and eosinophil counts (Fig. 7 and fig. S2). We believe this underlying asthmatic inflammation could have played a role in the exacerbated inflammatory response seen in *Scnn1b*-Tg mice during chronic infection. Finally, IL-17 is a key player in Th17-mediated (type 3) inflammation and T cell-mediated neutrophil mobilization and activation (*33, 42, 43*). IL-17-mediated inflammation was also described in CF patients and was correlated with pulmonary exacerbations and infection with *P. aeruginosa* (*44, 45*). The fact that KC/GRO and IL-17 were both upregulated during chronic infection in *Scnn1b*-Tg mice (Figs. 6 and 7, and Table S1) underscores the importance of the Th17/neutrophil interplay in the immune response of this model.

To counterbalance and resolve inflammation, regulatory T cells bear an important role by secreting the anti-inflammatory cytokine IL-10. Regulatory T cells were shown to be decreased in CF patients, and positively correlated with FEV1 (*66*). Compared to WT mice, regulatory T cells were increased in *Scnn1b*-Tg mice during infection (Fig. 5), which is opposite to what has been reported for CF patients. Nevertheless, their low proportion (1/10^4^ to 1/10^3^ T cells) (Fig. 5) and the almost undetectable levels of IL-10 (Fig. 6) suggest a limited role for the T cell subset in our model.

Another T cell subset present in our panel is the CD8^+^ T cell. Activated and central memory CD8^+^ T cells are important players of type 1 inflammation in response to intracellular pathogens and were upregulated in both WT and *Scnn1b*-Tg mice during chronic infection (Fig. 5). The marked upregulation in central memory CD8^+^ T cells (Fig. 5) and IL-15 secretion (Fig. 6) in WT mice following infection suggest a role for these cells in *P. aeruginosa* clearance. In *Scnn1b*-Tg mice, naïve CD8^+^ T cells were significantly increased, as for the CD8-derived double negative (DN) T cells (Fig. 5). Little is known about the role of these DN T cells during infection. Induction of DN T cells in murine models was described in response to intracellular pathogens such as *Leishmania major* and *Francisella tularensis* (*67–69*). In these studies, DN T cells were highly activated and produced IFNγ, TNFα, IL-17, and granzyme B. Although these DN T cells were protective against intracellular pathogens, it is unclear whether they had a role in *P. aeruginosa* clearance in our study. To our knowledge, elevated CD8^+^ and DN T cells are not described in CF patients thus, their presence during chronic infection in our mice may represent a limitation of using this model to study the inflammatory response to *P. aeruginosa* infection.

With its lung obstructive disease, underlying complex inflammation, tissue damage and inability to clear bacterial infection, we believe our SCFM-Tg-mouse model is a suitable model to study the host-pathogen interaction during chronic lung infections with *P. aeruginosa* and possibly with other pathogens. One primary limitation to our model is that it does not involve a CFTR deficiency, and thus cannot be used for modeling CFTR modulator therapies (*24, 70*). Because bacterial infections tend to persist in people with CF, even after CFTR modulator treatment (*71, 72*), it is important to consider alternative models for modulator-related studies, but also consider their limitations. Other rodent and non-rodent models have been used to reproduce the CF lung pathology (*9, 73*). CFTR-defective pigs and ferrets have similar lung pathologies to CF patients and are the only pre-clinical models to develop spontaneous lung infections (*74–79*). However, their severe intestinal disease, substantial cost and the strict legislations behind their usage in research can make these models challenging to use in research (*80*). Rat models including the CFTR knockout, the F508del CFTR and the humanized G551D models also develop defective in ion transport, airway mucus plugs, and multiorgan defects (*81–84*), making them appealing to study CFTR dysfunction using modulators. However, as for mice, rats do not develop spontaneous lung infection, and agar bead strategies were also used in this model to establish chronic infection (*85*). Furthermore, although these rats showed a higher neutrophilic response to infection (*84, 85*), it is not known whether they develop the complex asthmatic inflammation and lymphocytosis seen in CF patients and in our mouse model. Since our model utilizes SCFM2, and SCFM2 was shown to induce *P. aeruginosa* transcriptional profiles similar to those in human CF sputum in normal mice (*27*), we are confident that this SCFM-Tg-mouse model will be a valuable tool for investigating potential antimicrobials and the evolution of microbes during chronic infection.

## Materials and Methods

### Study Design

The objective of this study was to establish a chronic murine lung infection model in *Sccn1b-*Tg mice using SCFM2 agar beads laden with *P. aeruginosa* PAO1 and to determine the effects of this chronic infection on pulmonary function and inflammation. To determine these phenotypes, *P. aeruginosa* infected *Scnn1b-*Tg mice were compared to *P. aeruginosa* infected wild-type C57BL/6 mouse littermates, and sterile SCFM2 agar beads were also used as controls in both mouse genotypes to account for any possible bead specific effects. In addition, baseline analyses were performed on untreated *Scnn1b-*Tg mice and wild-type C57BL/6 mouse littermates. For all mouse experiments, 4-6 mice were used per group. Pulmonary function in each treatment group was measured as described below. Inflammation was determined by measuring immune cell populations by flow cytometry analyses, cytokine production, and lipid peroxidation. Mice were randomly assigned to groups, including equal distributions of males and females.

### Mouse model

B6N.Cg-Tg(Scgb1a1-Scnn1b)6608Bouc/J mice (*86*), herein named *Scnn1b-*Tg, were purchased from Jackson Laboratories (JAX stock #030949) and bred with C57BL/6 mice. Males and females 8-12 weeks old were equally distributed between the groups for experiments. *Scnn1b-*Tg mice were compared with their wild-type (WT) littermates. A total of 31 *Scnn1b-*Tg and 29 WT mice were used for experiments, divided into 4-6 mice/group. The Cedars-Sinai Institutional Animal Care and Use Committee approved all experiments according to current NIH guidelines.

### *P. aeruginosa* embedding in SCFM2-agar beads

*P. aeruginosa* PAO1 was obtained from Pradeep K. Singh (*87*) and grown in SCFM2 medium (*7, 8*). PAO1 embedding in SCFM2-agar beads was performed using a protocol adapted from Facchini *et al.* (*88*). Briefly, a single colony was inoculated into 3 mL SCFM2 and incubated at 37 °C overnight in a shaking incubator at 250 rpm. The next day, the culture was diluted in 7 mL of fresh SCFM2 and grown until a total of ∼5 OD was reached. In the meantime, 3% Bacto agar and 50mL of heavy mineral oil were autoclaved at 121°C for 45min and equilibrated at 50°C in a water bath. Bacto agar was then mixed with 2X SCFM2 pre-equilibrated at 50°C in a 1:1 ratio. Bacterial suspension was spun down, resuspended in 300 μL sterile PBS and mixed with 3mL of 1.5% Bacto agar 1X SCFM2 solution. The SCFM2 agar- *P. aeruginosa* mixture was added to heavy mineral oil and immediately stirred for 6 min at room temperature. The mixture was cooled to 4°C by stirring in iced water for 30 min. Agar beads were then transferred into 50 mL Falcon tubes and centrifuged at max speed for 15 min at 4 °C. Mineral oil was removed and agar beads were washed with sterile PBS 4 times. After the last wash, agar beads were resuspended in 25 ml PBS. To calculate the bacterial load of agar beads, an aliquot of the beads (approximately 0.5 ml) was aseptically homogenized and serially diluted 1:10 down to 10^-6^. Each dilution was spotted on LB plates and incubated at 37 °C overnight. The beads were stored at 4°C until the infection.

### Chronic infection

*Scnn1b-*Tg mice and WT littermates were anesthetized using 4% isoflurane. Sterile agar beads or 1×10^6^ CFU *P. aeruginosa*-laden SCFM2-agar beads were inoculated intratracheally using a 22-gauge angiocatheter (n=4-6 mice/ group). After 7 days of infection, either lung function measurements and CFU count, or flow cytometry were performed.

### Lung function measurements

The lung function was assessed by forced oscillation techniques and forced expiratory using the flexiVent FX system (SCIREQ)(*89*). The system was equipped with a FX2 module as well as with a NPFE extension for mice and it was operated by the flexiWare v7.2 software. Mice were anesthetized with isoflurane, intubated with a 18-20-gauge angiocatheter, and placed in the supine position in a plethysmograph chamber. Mice were mechanically ventilated at a tidal volume of 10mL/kg and frequency of 150 breath/min. The perturbations performed were a deep inflation, forced oscillation techniques (FOT), pressure-volume (PV) loop, and negative pressure-driven forced expiration (NPFE).

### CFU counts

After euthanasia, lungs were harvested and homogenized in sterile PBS using the Bead Mill 24 Homogenizer (Fisherbrand). The mixture was serially diluted 1:10 down to 10^-6^. Each dilution was spotted on LB plates and incubated at 37 °C overnight. CFUs were then counted and reported as CFUs/mL.

### Bronchoalveolar lavage for flow cytometry

Bronchoalveolar lavage (BAL) was performed with 6 x 1mL sterile cold 2mM EDTA/ 2% FBS/ 1X PBS buffer. BAL were spun at 500 rcf for 10 min at 4°C. Cell pellets were then resuspended in 3 mL RBC Lysis buffer and incubated at room temperature for 3 min. RBC lysis was stopped by adding 30 mL cold 3% FBS/ 1X PBS buffer. Cells were spun down, resuspended in 1.5 mL cold 3% FBS/ 1X PBS buffer, and counted using the TC20 Automated Cell Counter (Bio-Rad).

### Lung digestion for flow cytometry

Lungs were perfused through the right ventricle with 10 mL 1X PBS to flush blood out of lung tissue. Lungs were then removed, minced, and digested in 11mL 0.2% collagenase II (Worthington Biochem cat# LS004176) /10% FBS/RPMI 1640 media in a 37°C incubator shaking at 250rpm for 30 min. Digested lungs were then strained through 70 μm cell strainer and spun down at 500rcf for 10 min at 4°C. Cell pellets were then resuspended in 3 mL RBC Lysis buffer and incubated at room temperature for 3 min. RBC lysis was stopped by adding 30 mL cold 3% FBS/ 1X PBS buffer. Cells were spun down, resuspended in 5 mL cold 3% FBS/ 1X PBS buffer, and counted using the TC20 Automated Cell Counter (Bio-Rad). Cells were separated in different aliquots for inflammatory panel and lipid peroxidation assay.

### Inflammatory panel by flow cytometry

Up to 5×10^6^ cells were spun down in 1.5mL tubes at 8000 rcf for 1 min. Pellets were resuspended in 50 μL 3% FBS/ 1X PBS buffer with 2 μL FC block (BD cat# 553141) and incubated on ice for 20 min. Next, 50 μL of 3% FBS/ 1X PBS containing 0.25 μL of each cell surface antibody (see Table S2) was added and tubes were incubated on ice for 30 min in the dark. Cells were washed with 1 mL 3% FBS/ 1X PBS buffer and spun down. Cell pellets were then fixed in 500 μL cold 2% PFA and incubated at room temperature for 10 min with occasional vortexing to maintain single-cell suspension. Cells were spun down and washed with 1mL 3% FBS/ 1X PBS buffer. Cells were permeabilized in 150 μL 0.2% Tween-20/1X PBS buffer and incubated at room temperature for 15 min in the dark. Then, 50 μL of 0.2% Tween-20/1X PBS containing 1 μL of PE-Foxp3 (Miltenyi Biotec cat# 130-111-678) was added and cells were incubated for 30 min in the dark. Cells were washed with 1 mL 3% FBS/ 1X PBS buffer, spun down, and resuspended in 400 μL 1xPBS. Cell suspensions were filtered through a 70 μm mesh before analyzing on the Cytek Aurora spectral flow cytometer. Unmixing was performed with the Cytek SpectroFlo software version 3.1.0 and cell populations were analyzed on BD FlowJo version 10.8.2 and determined as follows (Figure 3): inflammatory cells (CD45^+^), neutrophils (Ly6G^+^), eosinophils (CD11b^+^, CD11c^-^, Siglec-F^+^), alveolar macrophages (Siglec-F^+^, CD11c^+^), classical monocytes (CD11b^+^, Ly6C^+^), monocyte-derived macrophages (CD11b^High^, Ly6C^+/-^, CD64^+^, FSC-A^high^), other myeloid-derived cells (CD11b^+^, Ly6C^-^), T cells (TCRβ^+^), T helper (TCRβ^+^, CD4^+^), Treg (TCRβ^+^, CD4^+^, Foxp3^+^), cytotoxic T cells (TCRβ^+^, CD8^+^), DN T cells (TCRβ^+^, CD4^-^, CD8^-^), naïve T cells (TCRβ^+^, CD44^-^, CD62L^+^), effector T cells (TCRβ^+^, CD44^+^, CD62L^-^), central memory T cells (TCRβ^+^, CD44^+^, CD62L^+^). The inflammatory antibody panel (Table S2) was designed using the EasyPanel V2 software (Omiq, LLC).

### Cytokine array

Total proteins were extracted from frozen lung tissue using Meso Scale Discovery MSD Tris Lysis Buffer (MSD cat# R60TX-3) supplemented with Protease Inhibitor Cocktail (Thermo Scientific cat# 78425), Phosphate Inhibitor Cocktail 2 (Sigma cat# P5726) and Phosphate Inhibitor Cocktail 2 (Sigma cat# P0044). Proteins were quantified using BCA Protein Assay (Genesee Scientific cat# 18-440). Cytokine array was performed using the V-PLEX Mouse Cytokine 29-Plex Kit (Meso Scale Discovery cat# K15267D-1). The following cytokines and chemokines were included: IFN-γ, IL-1β, IL-2, IL-4, IL-5, IL-6, IL-9, IL-10, IL-12p70, IL-15, IL-16, IL-17A, IL-17A/F, IL-17C, IL-17E/IL-25, IL-17F, IL-21, IL-22, IL-23, IL-27p28/IL-30, IL-31, IL-33, CXCL10, KC/GRO, MCP-1, MIP-1α, MIP-2, MIP-3α, TNF-α. The assay was performed following the manufacturer’s instruction and was analyzed on the Meso Scale Discovery instrument. Cytokine and chemokine levels were normalized to total proteins for the statistical analyses.

### Lipid peroxidation assay

Up to 5×10^6^ cells were spun down in 1.5mL tubes at 8000 rcf for 1 min. Pellets were resuspended in 100 μL of lipid peroxidation reagent 1:500 (Abcam cat# ab243377) and incubated for 30 min at 37°C and 5% CO_2_. Cells were washed with 1 mL 3% FBS/ 1X PBS buffer, spun down, and resuspended in 400 μL 1xPBS. Cell suspensions were filtered through a 70 μm mesh before analyzing on the BD Fortessa flow cytometer. Mean fluorescence intensity (MFI) was quantified using FlowJo software. Lipid peroxidation was quantified by calculating the red (Ex561/Em582)/green (Ex488/ Em525) fluorescence ratio. Data are presented as the reciprocal of the ratio (1/ratio).

### Statistical analysis

Normality and homogeneity of variance were assessed by Shapiro-Wilk and Brown-Forsythe tests, respectively. Data was log-transformed prior to analysis where necessary to meet assumptions necessary for parametric testing, else non-parametric rank testing was used. Based on data distributions, analyses between two groups were performed using Student t-test or the nonparametric Mann-Whitney. To detect any possible interaction between the mouse genotype and the infection on the parameters, ordinary two-way ANOVA followed by Tukey’s post-hoc test was used for comparisons between the four groups. Significant outliers determined by the Graph Pad Outlier Calculator were removed from statistics. All testing was considered significant at the two-tailed p-value of <0.05. Analysis performed with GraphPad Prism v10. The p-values are listed in Table S1.

## List of Supplementary Materials

Fig S1 to S3

Tables S1 to S2

## Acknowledgments

We would like to thank members of the Jorth Lab and Holly Huse for helpful discussions and feedback on this manuscript. We are grateful to Pradeep K. Singh for the generous gift of the P. aeruginosa strain used in this study. We also thank Christian Stehlik and Andrea Dorfleutner for the use of their Cytek Aurora flow cytometer. Finally, we are grateful to the Proteomics and Metabolomics Core at Cedars Sinai Medical Center for their help with the cytokine array analysis. This research was supported by grants from the Cystic Fibrosis Foundation to PJ (JORTH17F5, JORTH23I0, JORTH19P0) and the National Institutes of Health to PJ (R01AI14642) and a sub- award from UCLA CTSI National Institutes of Health grant UL1TR001881 to PJ.

## Author contributions include

Conceptualization: MV, PJ; Methodology: MV, PJ; Formal analysis: MV, DA, SEF, and PJ; Investigation: MV, DA, SEF, PJ; Resources: PJ; Data curation: MV, DA, SEF, PJ; Writing—original draft preparation: MV, PJ; Writing—review and editing: MV, DA, SEF, PJ; Supervision: MV, PJ; Project administration: MV, PJ; Funding acquisition: PJ.

